# Duplex real-time PCR assay for the simultaneous detection of *Achromobacter xylosoxidans* and *Achromobacter* spp

**DOI:** 10.1101/2020.02.11.944942

**Authors:** Erin P. Price, Valentina Soler Arango, Timothy J. Kidd, Tamieka A. Fraser, Thuy-Khanh Nguyen, Scott C. Bell, Derek S. Sarovich

## Abstract

Several members of the Gram-negative environmental bacterial genus, *Achromobacter*, are associated with serious infections in immunocompromised individuals, of which *Achromobacter xylosoxidans* is the most common. Despite their pathogenic potential, comparatively little is understood about these intrinsically drug-resistant bacteria and their role in disease, leading to suboptimal diagnosis and management of *Achromobacter* infections. Here, we performed comparative genomics of 158 *Achromobacter* spp. genomes to robustly identify species boundaries, to reassign several incorrectly speciated taxa, and to identify genetic sequences specific for the *Achromobacter* genus and for *A. xylosoxidans*. Next, we developed a Black Hole Quencher probe-based duplex real-time PCR assay, Ac-Ax, for the rapid and simultaneous detection of *Achromobacter* spp. and *A. xylosoxidans* from both purified colonies and polymicrobial clinical specimens. Ac-Ax was tested on 119 isolates identified as *Achromobacter* spp. using phenotypic or genotypic methods. In comparison to these routine diagnostic methods, the duplex assay showed superior identification of *Achromobacter* spp. and *A. xylosoxidans*, with five *Achromobacter* isolates failing to amplify with Ac-Ax confirmed to be different genera according to 16S rRNA gene sequencing. Ac-Ax quantified both *Achromobacter* spp. and *A. xylosoxidans* down to ∼110 genome equivalents, and detected down to ∼12 and ∼1 genome equivalent/s, respectively. *In silico* analysis, and laboratory testing of 34 non-*Achromobacter* isolates and 38 adult CF sputa, confirmed duplex assay specificity and sensitivity. We demonstrate that the Ac-Ax duplex assay provides a robust, sensitive, and cost-effective method for the simultaneous detection of all *Achromobacter* spp. and *A. xylosoxidans*, and will facilitate the rapid and accurate diagnosis of this important group of pathogens.

## Introduction

The *Achromobacter* genus comprises 19 officially designated species (1), all of which are highly motile, Gram-negative, non-fermentative bacteria found ubiquitously in environmental reservoirs including rivers, ponds, residential water sources, soil, mud, and some plants (2, 3). *Achromobacter* spp. are important community-acquired pathogens, particularly in people with cystic fibrosis (CF), cancer, immunoglobulin deficiencies, renal disease, endocarditis, diabetes, and those undergoing invasive procedures (4, 5). These opportunistic pathogens can infect several organs, although the respiratory and urinary tracts are the most common sites of infection (5). Members of this genus have been isolated from several usually sterile hospital products such as disinfectants, ultrasound gel, dialysis fluids, contact lens fluid, eardrops, incubators, respirators, humidifiers, and deionised water, consistent with the adaptability of *Achromobacter* spp. to survive in diverse environments (2, 4, 6).

Naturally multidrug-resistant bacteria, including *Achromobacter* spp., are increasingly being retrieved from CF airways due to the intensified implementation of aggressive antibiotic therapies (7, 8). *Achromobacter* spp. prevalence in CF centres globally range from 3 to 30%; of these, between 10 and 52% progress to a chronic infection. In addition to their intrinsic antibiotic resistance towards aminoglycosides, aztreonam, tetracyclines, and some penicillins and cephalosporins, *Achromobacter* spp. possess a similar denitrification system to *Pseudomonas aeruginosa*, which facilitates their survival and proliferation in hypoxic and anoxic environments such as those found in CF airways (5).

Although several *Achromobacter* species can infect CF airways (9), *Achromobacter xylosoxidans* is the most common, comprising ∼42-65% of all *Achromobacter* spp. identified in CF respiratory secretions (4, 10-12). Up until recently, the role of *Achromobacter* spp. in disease pathogenesis has been unclear; however, recent studies have shown that CF patients with an *Achromobacter* spp. infection are in fact at greater risk of experiencing a pulmonary exacerbation (9), and patients with chronic infections exhibit severe airway obstruction and more rapid lung function decline (13-15). Further, these pathogens can cause a range of serious diseases such as pneumonia, meningitis, osteomyelitis, urinary tract infections, and ocular infection in non-CF patients (16). Therefore, early and correct identification of *Achromobacter* spp. and *A. xylosoxidans* is important for improving the treatment and prognosis of disease.

Current diagnostic methods for identifying *Achromobacter* spp. and *A. xylosoxidans*, including the commonly used VITEK^®^ matrix-assisted laser desorption-ionisation time-of-flight mass spectrometry platform (VITEK^®^ MS), provide a reasonably accurate method for identifying these organisms (17). However, all *Achromobacter* spp. are allocated as *A. xylosoxidans*/*A. denitrificans* on this platform (18), thus providing limited capacity for accurate species-level identification. Further, VITEK^®^ MS and other mass spectrometry-based platforms require a purified isolate to obtain an accurate speciation result, which limits the utility of this platform as it cannot be used directly on polymicrobial clinical specimens such as sputum, resulting in longer turn-around times, potentially incorrect antimicrobial treatment, and higher costs (19, 20). In addition, mass spectrometry-based equipment has a large upfront cost and footprint, rendering this method out-of-reach for smaller, less-resourced laboratories. To address this shortcoming, an automated multiplex PCR has recently been developed to detect four non-fermentative Gram negative bacterial species, including *A. xylosoxidans*, directly from respiratory samples using the BD MAX™ System (18). This multiplex assay detected *A. xylosoxidans* with 97% specificity, but only 78% sensitivity (18), indicating suboptimal diagnosis of this organism using this method. Further, the BD MAX™ multiplex assay was not designed to identify other *Achromobacter* spp., meaning that ∼50% of CF infections caused by *Achromobacter* spp. cannot be diagnosed with this method. Other genotyping methods such as amplified ribosomal DNA restriction analysis (ARDRA) (21), multilocus sequence typing (22), *nrdA* gene sequencing (12), and whole-genome sequencing (WGS) provide robust identification and speciation methods for *Achromobacter* spp., but are laborious and cannot be performed in a rapid or cost-effective manner.

Here, we used a large-scale comparative genomic approach to identify genetic loci specific to all *Achromobacter* species, and to *A. xylosoxidans* only. We subsequently designed a highly specific and accurate *Achromobacter* spp. and *A. xylosoxidans* (Ac-Ax) duplex PCR assay for the simultaneous detection of these organisms. Phylogenomic analysis of 158 *Achromobacter* genomes, including 65 *A. xylosoxidans* genomes, was used to robustly identify species boundaries and to reassign several incorrect taxon assignments. Candidate genetic regions specific for all *Achromobacter* spp. and for *A. xylosoxidans* were then assessed for assay design suitability, followed by Ac-Ax duplex assay development and validation on 153 isolates comprising 116 *Achromobacter* spp., 48 *A. xylosoxidans*, and 34 non-*Achromobacter* spp. Finally, the Ac-Ax duplex assay was tested on 38 CF sputa DNA obtained from 21 adults, four of whom were positive for *Achromobacter* spp. according to 16S rRNA gene metataxonomic sequencing, to determine assay specificity and sensitivity in polymicrobial specimens.

## Materials and Methods

### *Achromobacter* spp. genomes and taxonomic reassignment

Publicly available data from 158 *Achromobacter* genomes was downloaded from the NCBI GenBank and Sequence Read Archive (SRA) databases (paired-end Illumina data only) in May 2019 (Table S1). Genomes from the following species were available for this study: *Achromobacter aegrifaciens* (*n*=1), *Achromobacter agilis* (*n*=1), *Achromobacter arsenitoxydans* (*n*=1), *Achromobacter denitrificans* (*n*=10), *Achromobacter dolens* (*n*=1), *Achromobacter insolitus* (*n*=8), *Achromobacter insuavis* (*n*=2), *Achromobacter marplatensis* (*n*=3), *Achromobacter mucicolens* (*n*=1), *Achromobacter piechaudii* (*n*=3), *Achromobacter pulmonis* (*n*=1), *Achromobacter ruhlandii* (*n*=7), *Achromobacter spanius* (*n*=6), *Achromobacter xylosoxidans* (*n*=79), and *Achromobacter* sp. (*n*=34). Following phylogenomic analysis of these data (described in the ‘Comparative genomics’ section below), taxonomic reassignment resulted in a final dataset comprising the following: *A. aegrifaciens* (*n*=3), *A. agilis* (*n*=1), *A. arsenitoxydans* (*n*=1), *A. denitrificans* (*n*=9), *A. dolens* (*n*=3), *A. insolitus* (*n*=9), *A. insuavis* (*n*=4), *A. marplatensis* (*n*=3), *A. mucicolens* (*n*=6), *A. piechaudii* (*n*=4), *A. pulmonis* (*n*=2), *A. ruhlandii* (*n*=17), *A. spanius* (*n*=5), *A. xylosoxidans* (*n*=65), and *Achromobacter* spp. (*n*=26) (Table S1).

Genome assemblies in GenBank that lacked corresponding raw reads in the SRA database were converted into simulated Illumina reads using ART version MountRainier (23). Prior to comparative genomic analysis, SRA data were quality-filtered by Trimmomatic v0.33 (24) using parameters described elsewhere (25).

### Comparative genomics to identify *Achromobacter* spp. and *A. xylosoxidans* loci

The methods for *in silico* identification of candidate conserved loci for *Achromobacter* spp. and *A. xylosoxidans* assay design have been detailed elsewhere (26, 27). Briefly, phylogenomic analysis was first performed to identify the *A. xylosoxidans* species boundary and to reassign incorrect species designations, followed by identification of conserved loci for the target taxa (i.e. all *Achromobacter* spp., and *A. xylosoxidans* only) among the 158 *Achromobacter* genomes (Table S1) using default parameters embedded in the SPANDx v3.2.1 comparative genomics software. The `-m` flag of SPANDx was employed to identify biallelic, orthologous, core-genome single-nucleotide polymorphisms (SNPs) among all genomes (28). Both simulated and real Illumina reads were mapped against the *A. xylosoxidans* NCTC 10807 reference genome (GenBank reference NZ_LN831029.1). Phylogenomic reconstruction was carried out on the 174,240 biallelic orthologous SNPs identified among all 158 *Achromobacter* spp. strains using the heuristic maximum parsimony function of PAUP* v4.0a.165 (29). The resultant phylogenomic tree (Figure 1) was bootstrapped for 1,000 replicates and midpoint-rooted using FigTree v1.4.0 prior to visualisation. The BEDcov output generated by BEDTools (30), which is wrapped in the SPANDx pipeline, was used to identify conserved candidate loci for subsequent real-time PCR assay design.

**Figure 1.**
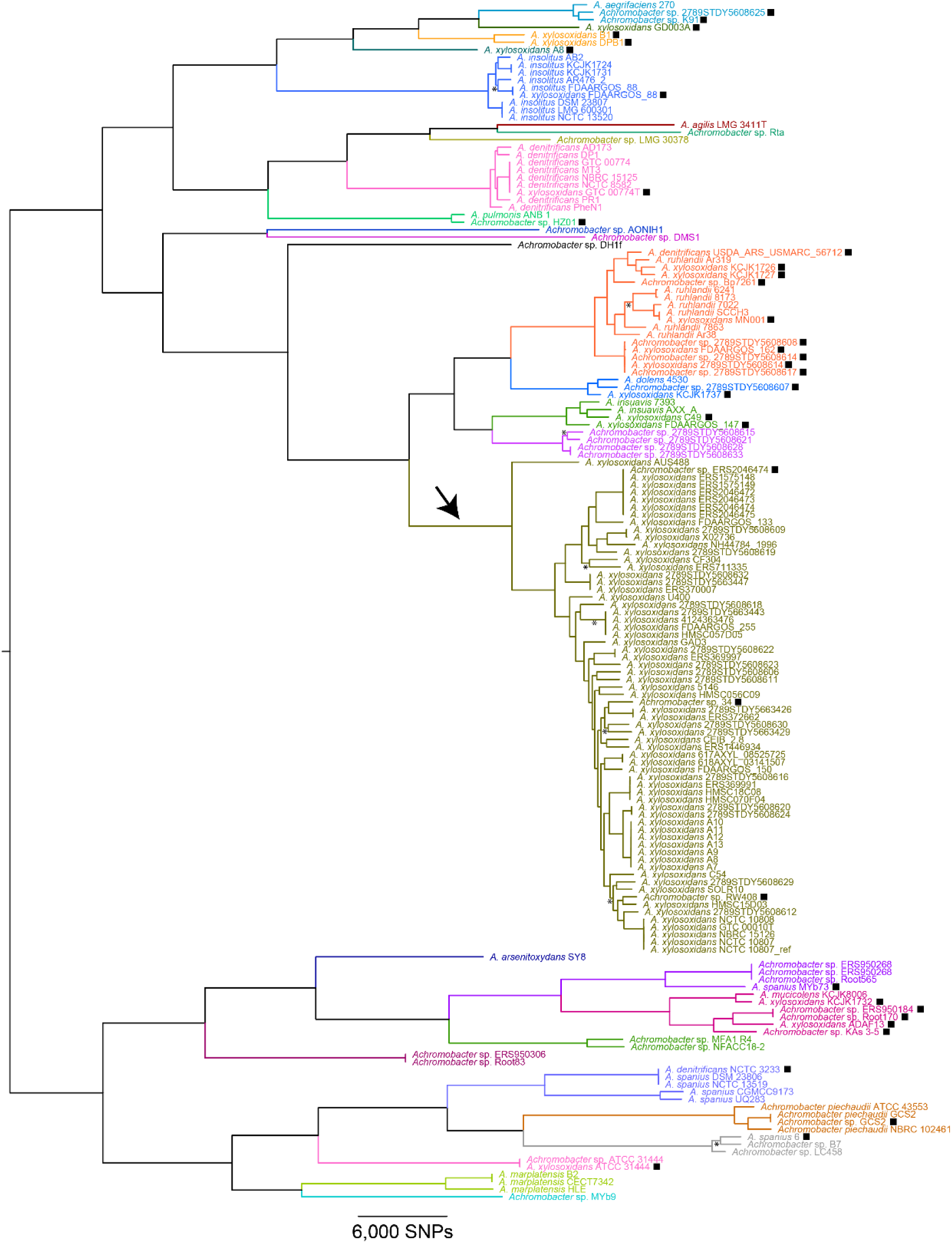
Maximum parsimony phylogenomic analysis of *Achromobacter* spp. This analysis reveals the large species diversity among *Achromobacter* spp., with >29 putative species identified within this genus based on publicly available genomic data. The delineation separating *A. xylosoxidans* from other *Achromobacter* spp. is shown by a black arrow. Incorrectly speciated taxa are shown by a black box next to the strain name. The most frequently misidentified species was *A. ruhlandii*, with 10 of the 17 *A. ruhlandii* strains incorrectly identified as *A. denitrificans* (*n*=1), *A. xylosoxidans* (*n*=5), or *Achromobacter* sp. (*n*=4). Branches with bootstrap values with <80% support are labelled by and asterisk. Consistency index=0.27.

### Identification of genetic loci specific for *A. xylosoxidans*, and for all *Achromobacter* spp

Using the BEDcov output generated by default by the SPANDx pipeline, a total of ∼9kb of DNA across four discrete loci was identified as highly conserved across all *A. xylosoxidans* strains (*n*=65), but absent or highly divergent in other *Achromobacter* spp. (*n*=94) (Table 1). The sequences for these loci were examined by Microbial Nucleotide BLAST (http://blast.ncbi.nlm.nih.gov; performed November 2019) to identify candidate regions for real-time PCR assay design. Using this approach, the *AT699_RS16685* locus, which encodes a hypothetical protein in *A. xylosoxidans*, was selected for assay design. The process for designing the *Achromobacter* spp. assay was different due to the need to cater for more genetic diversity across all *Achromobacter* strains. The highly conserved *rpoB* gene, which encodes DNA-directed RNA polymerase β-subunit protein, was targeted for assay design.

**Table 1.** Conserved loci in *Achromobacter xylosoxidans* (*n*=65 genomes) that are highly divergent or absent in other *Achromobacter* spp. (*n*=94 genomes) according to SPANDx.

| Genetic coordinates<br>(NCTC 10807*) | Encoded genes | Functions |
| --- | --- | --- |
| 1444000..1445000 | *AT699_RS06590* (partial),<br>*AT699_RS06595* (partial) | PAS domain-containing sensor histidine kinase, sensor histidine kinase |
| 3722000..3723000<sup>†</sup> | *AT699_RS16680* (partial),<br>*AT699_RS16685*,<br>*AT699_RS16690* | glutathione S-transferase family protein, hypothetical protein, winged helix-turn-helix transcriptional regulator |
| 4622000..4623000 | *AT699_RS20725* (partial),<br>*AT699_RS20730* | LysE family translocator, PhzF family phenazine biosynthesis protein |
| 5775000..5782000 | *AT699_RS26110* (partial),<br>*AT699_RS26115*,<br>*AT699_RS26120*,<br>*AT699_RS26125*,<br>*AT699_RS26130*,<br>*AT699_RS26135* (partial) | tripartite tricarboxylate transporter substrate binding protein, hypothetical protein, D-3-phosphoglycerate dehydrogenase, porin, long-chain-fatty-acid--CoA ligase, hypothetical protein |

\*GenBank reference NZ\_LN831029.1; <sup>†</sup>Locus targeted for PCR assay development in the current study.

### Ac-Ax duplex real-time PCR assay design

DNA sequences from these candidate loci for all 158 strains (for the Ac assay) and for the 65 *A. xylosoxidans* strains (for the Ax assay) were extensively assessed for specificity using several *in silico* methods as detailed elsewhere (31). The following primers and Black Hole Quencher (BHQ) probes were designed for specific *Achromobacter* spp. and *A. xylosoxidans* detection, respectively (5’ to 3’): Ac_F (CACrTAGCTCACGAACTCCAAGC), Ac_R (CAGCTTCAATCCTACCTAACTTTCCT), and Ac_probe (HEX-CGTAGCCGACGGTTTGCAGG-BHQ1), which generates a 144bp amplicon; and Ax_F (AGCGTCACGGAATGCAGC), Ax_R (AAGGGCGTTTCAACGAGAGC) and Ax_probe (FAM-AGGTCATAGGCGTAGACCAGC-BHQ1), which generates a 127bp amplicon. Amplicon specificity was verified using Microbial Nucleotide BLAST.

### Real-time PCR parameters

PCR optimisation was performed for both assays in singleplex across a range of primer (0.2, 0.25, 0.3, 0.35, and 0.4 µM) and subsequently probe (0.25, 0.3, 0.35, 0.4 µM) concentrations to determine the optimal oligomer concentrations for each assay prior to conversion to the duplex format. The optimised Ac-Ax duplex PCR consisted of 1X Sso Advanced Universal Probes Supermix (Bio-Rad Laboratories, Gladesville, NSW, Australia), 0.40 µM of the Ax_probe, Ac_F, and Ac_R oligomers, 0.35 µM Ac_probe, and 0.25 µM Ax_F and Ax_R oligomers (Macrogen Inc., Geumcheon-gu, Seoul, Rep. of Korea), 1 µL DNA template, and RNase/DNase-free PCR grade water (Thermo Fisher Scientific), to a final volume of 5 µL reaction volume. Thermocycling was performed using the CFX96 Touch™ Real-Time PCR Detection System (Bio-Rad), with parameters consisting of an initial hot start/denaturation step of 95 °C for 2 min, followed by 45 cycles of denaturation at 95 °C for 5 sec and annealing/extension at 60 °C for 5 sec. The *A. xylosoxidans* LMG 1863 type culture strain (32) was used as a control for all experiments, with no-template controls (NTCs) included in all runs to assess assay performance. All PCR results were examined using the CFX Maestro v4.1.2433.1219 software (Bio-Rad).

### Analysis of *Achromobacter* spp. strains using the Ac-Ax PCR assay

We examined the performance of our duplex assay across *A. dolens* LMG 26840 and *A. insuavis* LMG 26845 (33), *A. ruhlandii* LMG 1866 (34), *A. xylosoxidans* LMG 1863 (32), and 115 Australian strains identified as *Achromobacter* spp. according to: i) VITEK^®^ MS microbiological testing (*n*=12), ii) API 20 NE phenotypic testing (*n*=85); iii) ARDRA (*n*=13), and iv) WGS (*n*=5) (Table S2). Among the Australian strains, two were previously identified as *A. ruhlandii* (QLDACH007 and QLDACH010) and two as *A. xylosoxidans* (QLDACH001 and AUS488) according to WGS (35, 36), one (QLDACH016) was a novel *Achromobacter* sp. according to WGS (36), and 110 were allocated as *Achromobacter* sp. according to API 20 NE, ARDRA, or VITEK^®^ MS. Strains were grown on chocolate agar for 24h at 37 °C prior to chelex DNA extraction, as described elsewhere (31).

### Ac-Ax PCR assay sensitivity and specificity testing

To determine Ac-Ax PCR assay sensitivity, the limits of detection (LoD) and quantification (LoQ) were determined (31) in the duplex assay format using serial dilutions of *A. xylosoxidans* LMG 1863 DNA ranging from 40 ng/µL to 0.04 fg/µL. Twenty-four no-template controls (NTCs) were also included. Next, the Ac-Ax duplex assay was tested for specificity against 34 non-*Achromobacter* isolates comprising *Burkholderia* spp. (*n*=3), *Enterobacter* spp. (*n*=4), *Klebsiella* spp. (*n*=3), *Prevotella* spp. (*n*=4), *P. aeruginosa* (*n*=9), *Staphylococcus* spp. (*n*=3), *Stenotrophomonas maltophilia* (*n*=1), and *Veillonella* spp. (*n*=6). These organisms were selected as they represent a cross-section of species identified in human infections, particularly in CF airways.

### 16S rRNA gene sequencing

Thirty-eight sputa from 21 adults with CF presenting at a single CF clinic in Brisbane, Australia, were subjected to 300bp paired-end Illumina MiSeq 16S rRNA gene metataxonomic sequencing to identify the presence of achromobacterial DNA in these polymicrobial specimens. These data were compared with the Ac-Ax assay to determine the performance of this duplex PCR on polymicrobial specimens. The V3-V4 region was targeted using the universal 341F (5’-CCTAYGGGRBGCASCAG) and 806R (5’-GGACTACNNGGGTATCTAAT) primers, with PCRs, sequencing, and data analysis performed at the Australian Genome Research Facility (St Lucia, Qld, Australia) according to standardised workflows.

16S rRNA gene sequencing was carried out on five *Achromobacter* isolates (as determined by ARDRA or API 20 NE testing) that were negative for the Ac-Ax duplex assay to assign species designations. The ∼1.3kb 16S rDNA amplicons were generated using primers 785F (5’-GGATTAGATACCCTGGTA) and 907R (CCGTCAATTCMTTTRAGTTT), followed by dideoxy sequencing at Macrogen Inc. (Geumcheon-gu, Seoul, Republic of Korea). Sequence chromatograms were visualised in BioEdit v7.2 (37).

### 16S rRNA gene universal real-time PCR

A universal bacterial 16S rRNA gene SYBR Green assay comprising oligos 16S-UniF (5’-TCCTACGGGAGGCAGCAGT) and 16S-UniR (5’-GGACTACCAGGGTATCTAATCCTGTT) (38) was used for relative bacterial DNA quantitation across the 38 CF sputa. Reactions were carried out in duplicate in a final 5 µL volume using the same mastermix, real-time PCR instrumentation, and DNA volumes as described above for the Ac-Ax assay. Minor modifications were made to the thermocycling parameters as follows: initial denaturation for 2 min at 95°C, followed by 40 cycles of denaturation for 15 sec at 95°C, and 20 sec annealing/extension at 60°C. The relative abundance of achromobacterial DNA in these CF sputa was determined by subtracting the *Achromobacter* sp. cycles-to-threshold (C_T_) value (where positive) from the 16S rRNA gene C_T_ value (i.e. ΔC_T_).

## Results

### Comparative genomic analysis of *Achromobacter* spp

Phylogenomic reconstruction of the 158 *Achromobacter* genomes confirmed that all taxa were *Achromobacter* spp. However, a considerable number of taxonomic errors (*n*=36; ∼23%) were identified in the dataset (Figure 1, black boxes). Taxonomic reassignment was therefore carried out to ensure correct delineation of the *A. xylosoxidans* clade from all other *Achromobacter* spp. for PCR assay design (Table S1). In total, 29 *Achromobacter* clades were identified among the 158 genomes, of which only 14 corresponded to a previously assigned species. The remaining 15 clades lack a type species genome for comparison and may correspond to several novel species (Figure 1).

### BLAST analysis identifies additional *Achromobacter* spp. and *A. xylosoxidans* misclassifications

Microbial Nucleotide BLAST analysis of the Ac-Ax duplex assay amplicons against 31 complete and 141 draft *Achromobacter* spp. genomes, and 32,972 complete non-*Achromobacter* and 9,040 draft Betaproteobacterial genomes (as at 25-Nov-19) identified a small number strain misclassifications in the NCBI database. BLAST analysis of the 144bp Ac amplicon identified one putative false-positive hit (*Bordetella bronchiseptica* strain KU1201; contig BBVB01000043.1); however, closer inspection showed greater homology of this contig to *Achromobacter* spp. (∼95-99% identity and 97-100% coverage) than *Bordetella* spp. (∼90-91% identity and ∼68-71% coverage). The closest non-*Achromobacter* hit for the Ac amplicon was in *Bordetella* genomosp. 7 (∼89% identity and 100% coverage). Importantly, there were 6 SNPs in the 20bp Ac_probe sequence in these taxa, which would inhibit their detection in the real-time PCR assay due to insufficient sequence homology. For all 172 *Achromobacter* spp. genomes, there was 100% nucleotide conservation at the primer- and probe-binding regions. Therefore, *in silico* analysis confirmed excellent specificity of the Ac assay for all known members of this genus.

For the 127bp Ax amplicon, one putative false-positive BLAST hit was identified (*Achromobacter* sp. RW408); however, this isolate was reclassified as *A. xylosoxidans* according to our phylogenomic analysis (Figure 1), confirming an NCBI database error for this strain. The closest non-*A. xylosoxidans* hit was in *Burkholderia mesoacidophila*, with BLAST analysis yielding 100% coverage but only 75% sequence identity in this organism. Like the Ac assay, there were 6 SNPs in the 21bp Ax_probe sequence in *B. mesoacidophila*, which would inhibit detection of this non-target species due to substantial sequence diversity. All 55 *A. xylosoxidans* genomes possessed 100% nucleotide conservation at the primer- and probe-binding sites. Therefore, this assay shows excellent *in silico* specificity for *A. xylosoxidans*.

### Ac-Ax performance on *Achromobacter* isolates

Of the 119 *Achromobacter* isolates examined with the Ac-Ax duplex real-time PCR assay, 114 were Ac-positive, and among these, 48 were also Ax-positive (Table S2). There were no instances of Ax-positive but Ac-negative strains. The four type culture strains performed as expected, with all being Ac-positive, and only *A. xylosoxidans* LMG 1863 being Ax-positive; *A. insuavis* LMG 26845, *A. ruhlandii* LMG 1866, and *A. dolens* LMG 26840 failed to amplify with the Ax assay. The five isolates that did not amplify with either assay were subjected to 16S rRNA gene sequencing of a ∼1.3kb amplicon to determine their species identity. Of these, two (QLDACH029 and QLDACH035) were identified as *P. aeruginosa*, and the remaining three were identified as *B. bronchiseptica* (QLDACH105), *Cupriavidus metallidurans* (QLDACH120), and *S. maltophilia* (QLDACH125). The two *P. aeruginosa* isolates were previously identified as *Achromobacter* sp. according to ARDRA (39), whereas the remaining three isolates were identified as *Achromobacter* sp. according to API 20 NE. The performance of each genotyping method and concordance with the Ac-Ax duplex assay is summarised in Table 2.

**Table 2.** Summary of genotyping methods for *Achromobacter* identification and performance comparison with the Ac-Ax duplex real-time PCR assay.

| Initial ID method | No. Ac-positive (% concordance) | No. Ax-positive (% concordance) |
| --- | --- | --- |
| WGS (*n*=5) | 5 (100%) | 2 (100%) |
| ARDRA (*n*=13) | 11 (84.6%) | N/A |
| VITEK® MS (*n*=12) | 12 (100%) | N/A |
| API 20 NE (*n*=85) | 82 (96.5%) | N/A |

Abbreviations: Ac, *Achromobacter* sp.; ARDRA, amplified rRNA gene restriction analysis; Ax, *Achromobacter xylosoxidans*; MS, mass spectrometry; N/A, not applicable (due to lack of resolution); WGS, whole-genome sequencing

### Ac-Ax performance on non-*Achromobacter* isolates

Of the 34 non-*Achromobacter* species and 24 NTCs tested against the Ac-Ax duplex assay, none yielded detectable amplification (data not shown).

### Ac-Ax sensitivity

The lower limits of detection (LoD) and quantification (LoQ) for the Ac-Ax duplex assay were determined on *A. xylosoxidans* LMG 1863 genomic DNA obtained from a pure culture (Figure 2). Using a 10-fold DNA dilution series ranging from 40 ng/µL to 0.04 fg/µL, the LoQ for both assays was ∼400 fg/µL, or ∼110 genome equivalents (GEs). The LoD values were more sensitive than the LoQ values, with an Ac assay LoD of ∼40 fg/µL (∼12 GEs) and an Ax LoD of ∼4 fg/µL (∼1 GE) (Figure 2).

**Figure 2.**
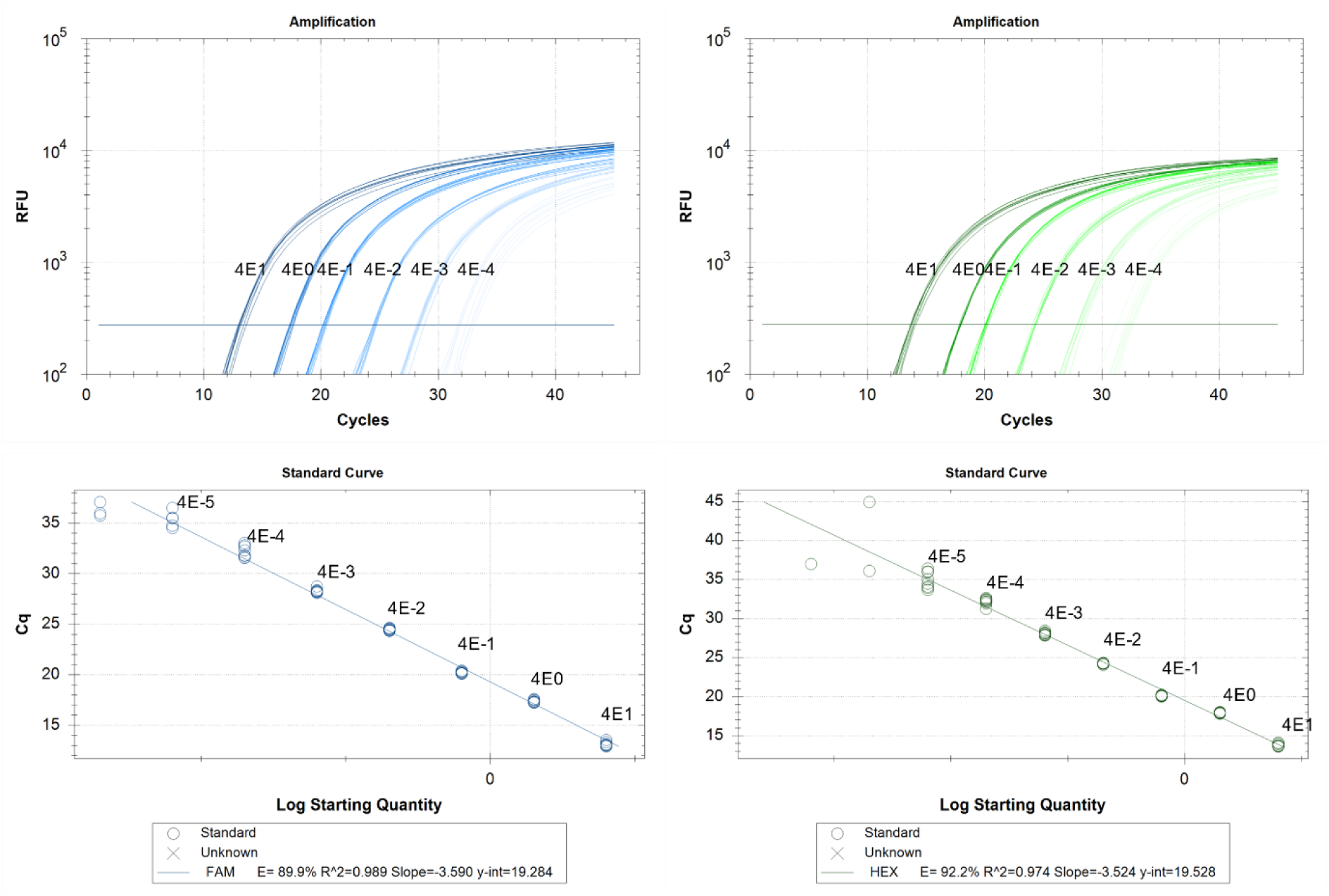
Limits of detection (LoD) and quantitation (LoQ) for the *Achromobacter* spp. (Ac; left) and *Achromobacter xylosoxidans* (Ax; right) duplex real-time PCR assay across a standard curve. Both assays yielded LoQ values of ∼400 fg/µL (∼110 genome equivalents). The Ac assay exhibited an LoD value of ∼40 fg/µL (∼12 genome equivalents), whereas the Ax assay yield an LoD value of ∼4 fg/µL (∼1 genome equivalent).

### Comparison of Ac-Ax and metataxonomics for *Achromobacter* identification from CF sputa

To determine the performance of Ac-Ax on polymicrobial clinical specimens, the duplex PCR was tested against 38 sputa from 21 adults with CF. Of these, five (i.e. 15%) contained *Achromobacter* spp. at relative abundances ranging from 0.1% to 63.6% according to metataxonomic sequencing (Table 3), with the remaining 33 samples failing to identify any *Achromobacter* 16S rRNA gene reads. Consistent with the metataxonomic findings, 4/33 sputa were PCR-positive according to the Ac assay, and three of these were also Ax-positive; however, this species result could not be compared with the metataxonomic data due to insufficient species-level resolution obtained from the 16S rRNA gene V3-V4 region.

**Table 3.** Performance comparison of the Ac-Ax real-time duplex PCR assay against 16S rDNA metataxonomic sequencing on five *Achromobacter*-positive sputa obtained from cystic fibrosis airways.

| Sample | C<sub>T</sub> | | $\Delta C_T^*$ | 16S rDNA metataxonomic sequencing (% relative abundance) |
| --- | --- | --- | --- | --- |
|  | Ac | Ax |  |  |
| *A. xylosoxidans* LMG 1863 | 18.4 | 18.5 | ND | N/A |
| SCHI0014 | 36.3 | Neg | 15.2 | *Achromobacter* sp. (0.1%) |
| SCHI0003 | 28.3 | 26.0 | 4.5 | *Achromobacter* sp. (35.5%) |
| SCHI0009 | 25.0 | 25.5 | 2.8 | *Achromobacter* sp. (63.6%) |
| SCHI0030 Day 1 | 34.7 | 34.7 | 9.3 | *Achromobacter* sp. (0.4%) |
| SCHI0030 Day 6 | Neg | Neg | N/A | *Achromobacter* sp. (0.1%) |

\*Determined by subtracting the Ac cycles-to-threshold (C<sub>T</sub>) from the 16S rDNA C<sub>T</sub>. Abbreviations. Ac, *Achromobacter* sp.; Ax, *A. xylosoxidans*; N/A, not applicable; ND, not determined; Neg, negative PCR result

The relative abundance of achromobacterial DNA between the metataxonomic and duplex PCR methods was also consistent. For example, the highest proportion of achromobacterial DNA was detected in SCHI0009 (difference in cycles-to-threshold [ΔC_T_] value of 2.8 when compared with the 16S rRNA gene), which possessed the highest relative abundance of achromobacterial reads (63.6%) according to metataxonomics (Table 3). The one Ac-Ax negative sample, SCHI0030 Day 6, only contained a relative abundance of 0.1% achromobacterial DNA in the metataxonomic sequence data. Sputa from another patient, SCHI0014, had the same low relative abundance of achromobacterial DNA but was Ac-positive; however, the C_T_ value (36.3) was found to be outside the LoQ and LoD values for this assay (Figure 2), thus reflecting the stochastic nature of detection capability beyond these limits. The higher sensitivity of the metataxonomic method for achromobacterial detection was also expected due to the multicopy nature of the 16S rRNA gene (*n*=3) in *Achromobacter* spp. compared with the single-copy nature of the Ac and Ax targets.

## Discussion

*Achromobacter* spp. are now recognised as an important cause of severe nosocomial and community-acquired infections. These Gram-negative bacteria can cause a spectrum of disease including bacteraemia, cholecystitis, endocarditis, keratitis, lymphadenitis, meningitis, osteomyelitis, peritonitis, pneumonia, and urinary tract infections (40, 41). *Achromobacter* spp. are becoming increasingly common in people with the life-shortening disease, CF, being present in up to 30% of adult CF airways (13, 15, 42). Although historically considered of low pathogenic potential, there is mounting evidence that CF infections caused by *Achromobacter* spp. are associated with adverse clinical presentations and outcomes, especially in immunocompromised individuals (9, 13-15). Therefore, their rapid identification is essential for guiding appropriate therapeutic treatments and improving patient prognosis (43).

Several phenotypic (e.g. API 20 NE, VITEK^®^ MS) and genotypic (e.g. ARDRA, gene sequencing, WGS, real-time PCR) methods are available to identify *Achromobacter* spp. These methods provide varying degrees of sensitivity, specificity, cost-effectiveness, turnaround time, and resolution. The gold standard method, next-generation sequencing, enables highly accurate and comprehensive species identification, but is currently laborious, slow (>8h to result), costly (∼AUD$80 for metataxonomics or WGS), and requires specialised bioinformatic tools and knowledge to analyse sequence data. VITEK^®^ MS is considered to have good success at identifying *Achromobacter* spp. to the genus level; however, this method currently cannot attain reliable species-level resolution, with e.g. *A. xylosoxidans* unable to be differentiated from *A. denitrificans* (18), despite these species being genetically distinct (Figure 1).

We chose the real-time PCR platform for Ac-Ax assay development due to its multiplexing capability, low per-sample cost, high accuracy potential, direct detection from polymicrobial specimens (e.g. sputum), excellent sensitivity, greater accessibility in lower-resourced laboratories, and rapid (<1h) turnaround time (18). The upfront equipment cost of real-time PCR equipment (∼USD$25-40K) is also considerably less than VITEK^®^ MS (USD$200K) or many next-generation sequencing platforms such as Illumina, and has a much smaller laboratory footprint. The Ac-Ax consumables cost is comparable to VITEK^®^ MS at ∼USD$1 per sample, compared with ∼USD$30 for the BD Max real-time PCR platform, making the Ac-Ax assay a cost-effective method for achromobacterial identification. The Ac-Ax assay also has the advantage of simultaneous detection of both *Achromobacter* sp. and *A. xylosoxidans*, unlike most existing methods that only detect *A. xylosoxidans*, meaning that ∼50% of *Achromobacter* CF infections are undiagnosed with these methods (4, 10-12).

The Ac-Ax assay is highly accurate, with no false-positives or false-negatives identified with *in silico* or laboratory testing. Indeed, our initial *in silico* BLAST analysis of Ac and Ax targets resolved incorrect species assignments in two publicly available genomes: *B. bronchiseptica* KU1201 (actually *Achromobacter* sp.) and *Achromobacter* sp. RW408 (actually *A. xylosoxidans*), demonstrating the highly accurate nature of these targets. Laboratory testing of the Ac-Ax duplex assay identified 114 of 119 previously characterised *Achromobacter* isolates as *Achromobacter* spp., of which 48 (42%) were *A. xylosoxidans* (Table S2). The five Ac-Ax negative isolates, which were incorrectly identified as *Achromobacter* sp. according to API 20 NE or ARDRA, were confirmed as *B. bronchiseptica, C. metallidurans, P. aeruginosa* or *S. maltophilia* based on 1.3kb 16S rRNA gene sequencing. All five genus misclassifications were also confirmed by WGS (data not shown). Previous studies have demonstrated poor performance of API 20 NE for *Achromobacter* spp. identification, with *Bordetella petrii, Bordetella trematum, Ralstonia pickettii, Alcaligenes faecalis, Comamonas testosteronii, Moraxella* sp., *Pasteurella* sp., and *Pseudomonas alcaligenes* being incorrectly called as *Achromobacter* spp., or vice versa (44-47). Based on these collective findings, we do not recommend ARDRA or API 20 NE for *Achromobacter* identification due to relatively high false-positivity rates with other common pathogens.

A major component of highly robust microbial diagnostics development is ensuring correct species identification and locus specificity for the target taxa. The gold standard method for achieving high-quality target identification is large-scale comparative genomic analysis to identify species- and genus-level boundaries. As demonstrated by our phylogenomic analysis (Figure 1), nearly a quarter of the 158 publicly deposited *Achromobacter* genomes were incorrectly speciated. Our study therefore demonstrates the critical importance of using large-scale comparative genomics for informed and accurate diagnostics development.

The Ac-Ax assay demonstrated good sensitivity and specificity for achromobacterial identification from polymicrobial specimens, with 4/33 (12%) adult CF sputa being Ac PCR-positive. This rate falls within the 3 to 30% *Achromobacter* prevalence rates reported in CF centres worldwide (13, 15, 42). Of the four positive sputa, one patient was found to harbour a non-*A. xylosoxidans* achromobacterial infection (Table 3). The Ac-Ax results were consistent with metataxonomic sequencing, which identified 5/33 (15%) achromobacterial-positive sputa. The one Ac-Ax PCR-negative sputum sample, collected of Day 6 of intravenous antibiotic treatment in Patient SCHI0030, had a very low (∼0.1%) proportion of achromobacterial rDNA gene reads that exceeded the lower Ac-Ax LoD threshold (∼12 and ∼1 genome equivalent/s for Ac and Ax, respectively). Notably, the Ac-Ax assay detected very low prevalence of *A. xylosoxidans* in the Day 1 sputum sample from Patient SCHI0030 (C_T_=34.7 for both Ac and Ax assays), which corresponded with a similarly low (∼0.4%) proportion of achromobacterial rRNA gene reads, indicating very low prevalence of this organism in this patient’s airways at both time points. Taken together, we show that the Ac-Ax assay provides good performance on polymicrobial specimens, with the advantage of a considerable reduction in cost and turnaround-time compared with metataxonomic sequencing.

In conclusion, we have employed a large-scale comparative genomics approach to inform the design of a highly accurate duplex real-time PCR assay for the rapid, sensitive, specific, cost-effective, and simultaneous detection of *Achromobacter* sp. and *A. xylosoxidans* from purified cultures and polymicrobial clinical specimens. Implementation of the Ac-Ax assay in the clinic will enable rapid (∼1h) diagnosis of these naturally drug-resistant organisms, providing the opportunity for targeted antimicrobial therapy and rapid treatment shifts in response to achromobacterial detection. Although beyond the scope of the current study, future work should compare Ac-Ax and VITEK^®^ MS results across a large isolate panel to determine VITEK^®^ MS accuracy among *Achromobacter* spp.

## Author Statements

### Authors and Contributions

Conceptualization: EPP, DSS; methodology and experimental design: EPP, VSA, TAF, DSS; sample collection: TJK, T-KN, SCB; original draft preparation: EPP, VSA; manuscript review and editing: TJK, TAF, T-KN, SCB, DSS.

### Ethics approval

This study was approved by The Prince Charles Hospital Human Research Ethics Committee (HREC/13/QPCH/127).

## Acknowledgements

We wish to thank Timothy Wells and Amy Pham (Translational Research Institute, Brisbane, Australia) for providing *Prevotella* and *Veillonella* DNA for this study.

## Supplemental data

**Table S1.**
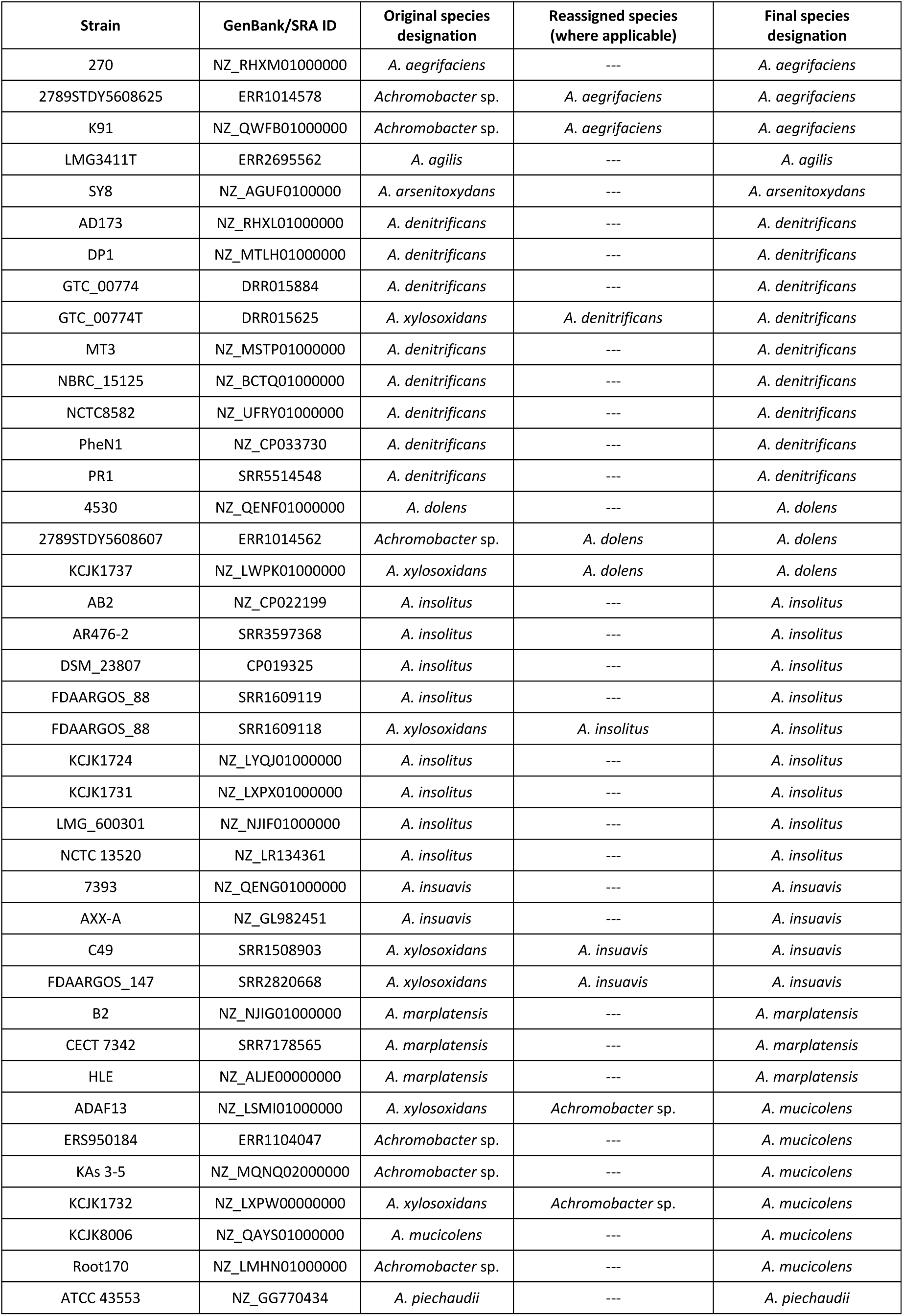
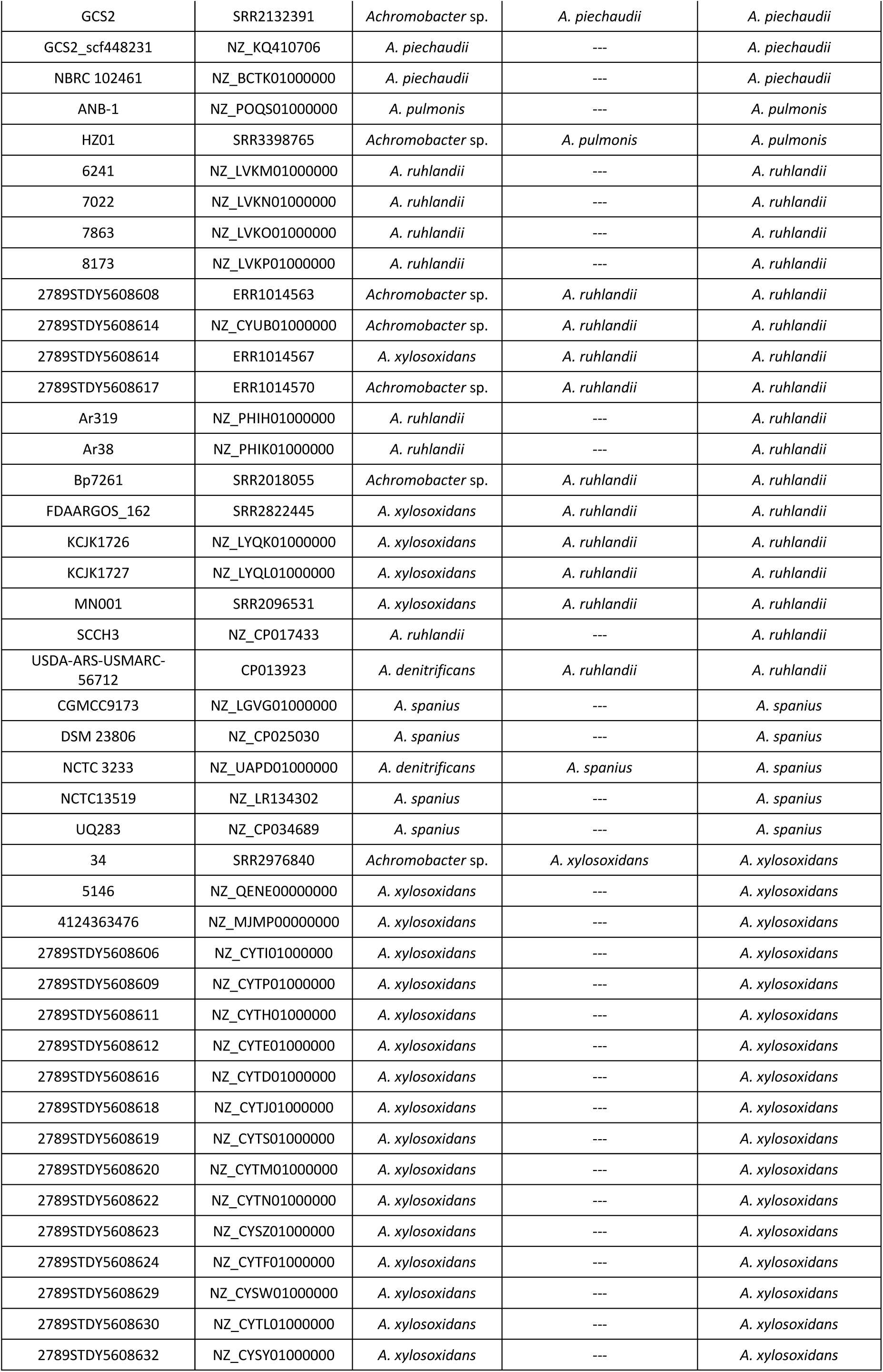
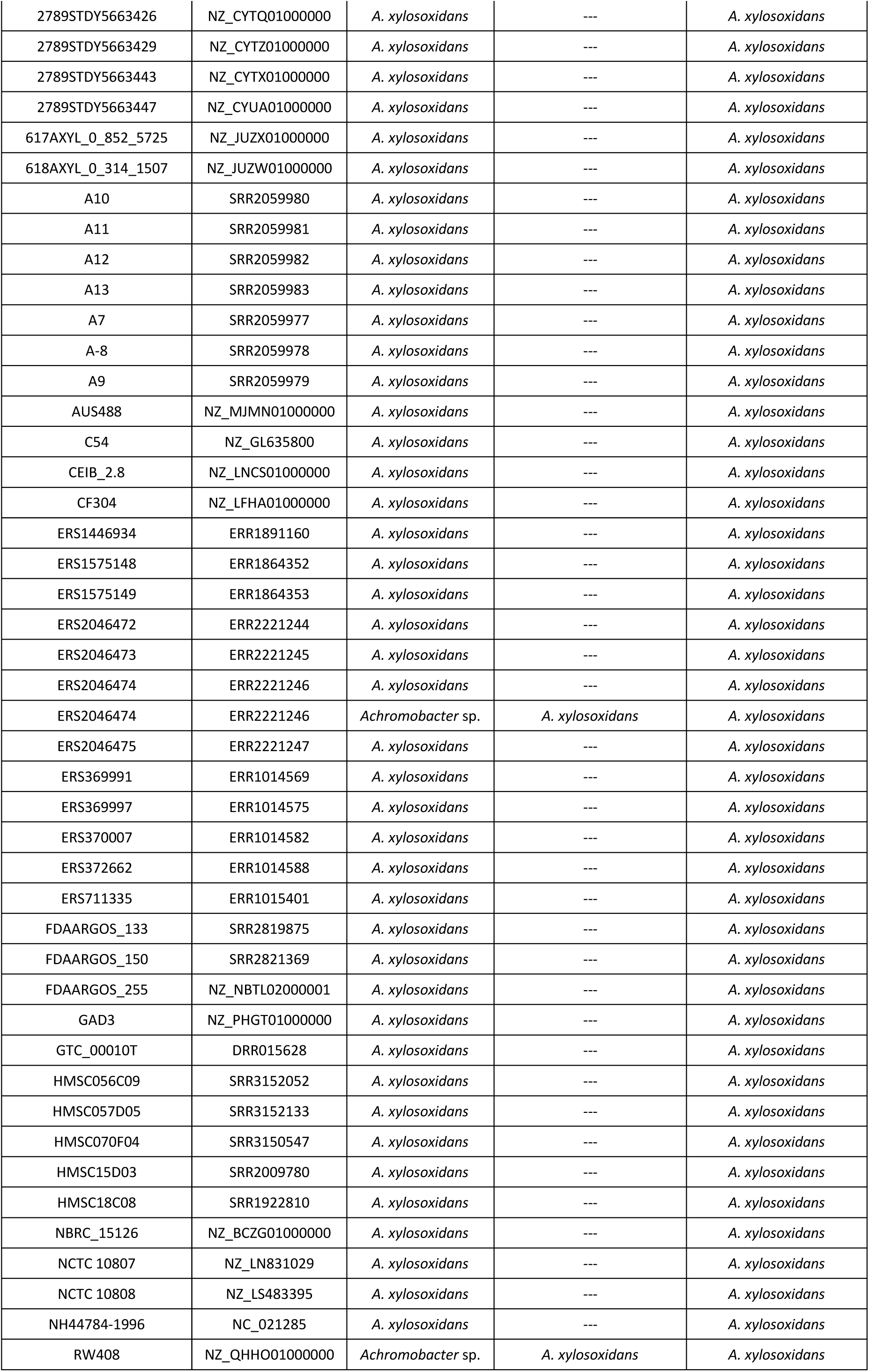
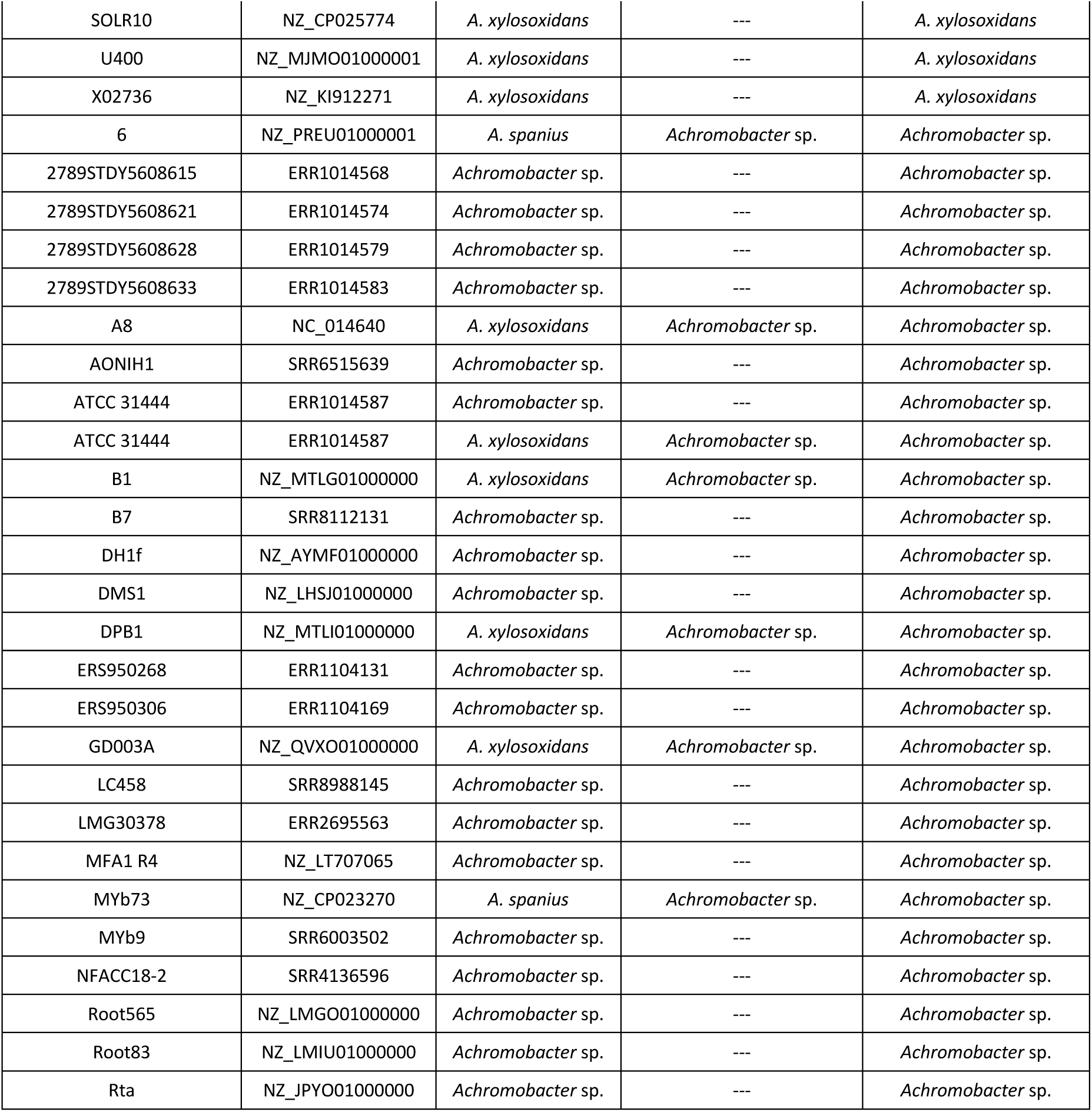
List of the 158 *Achromobacter* genomes used in this study.

**Table S2.**
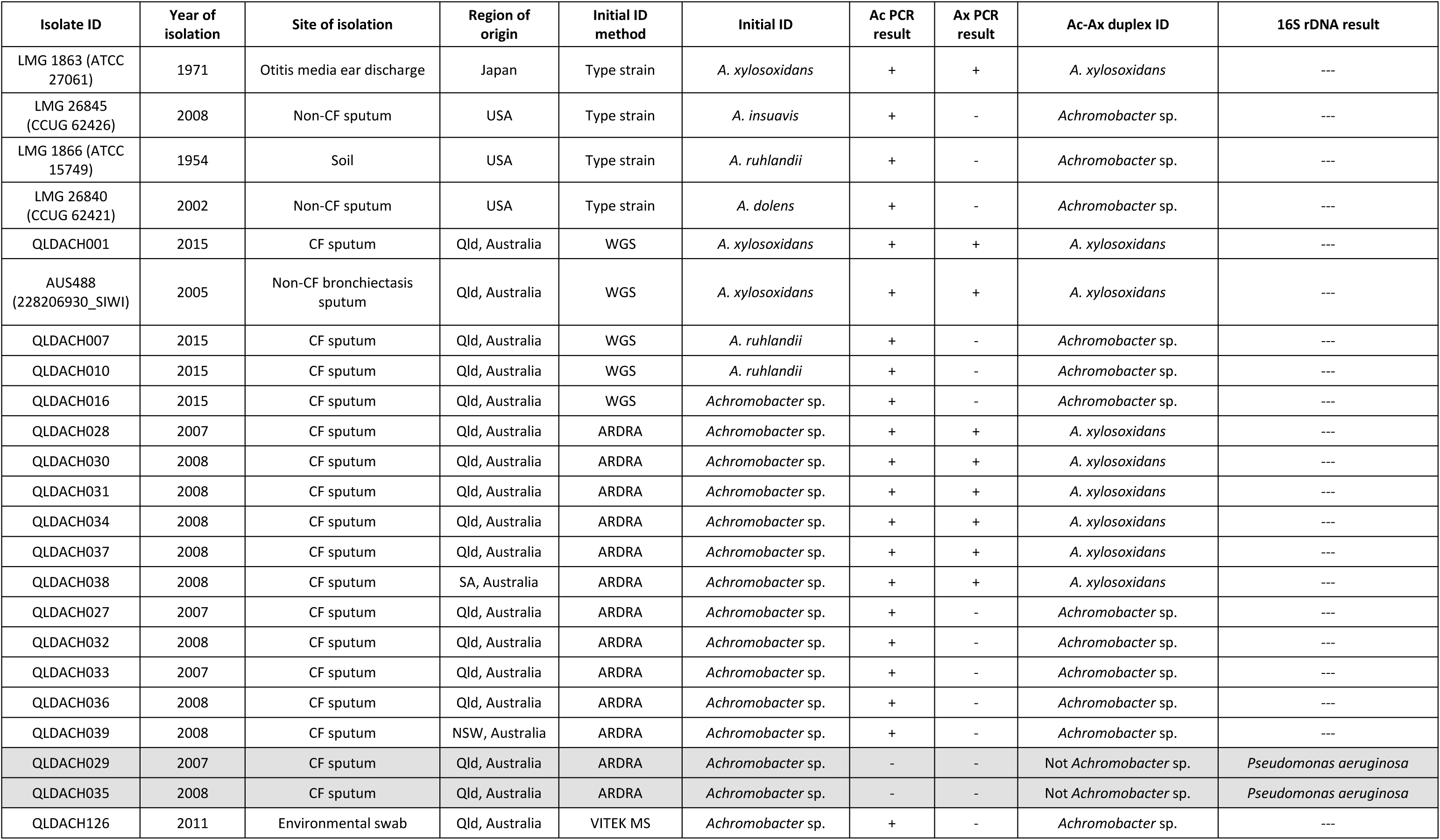
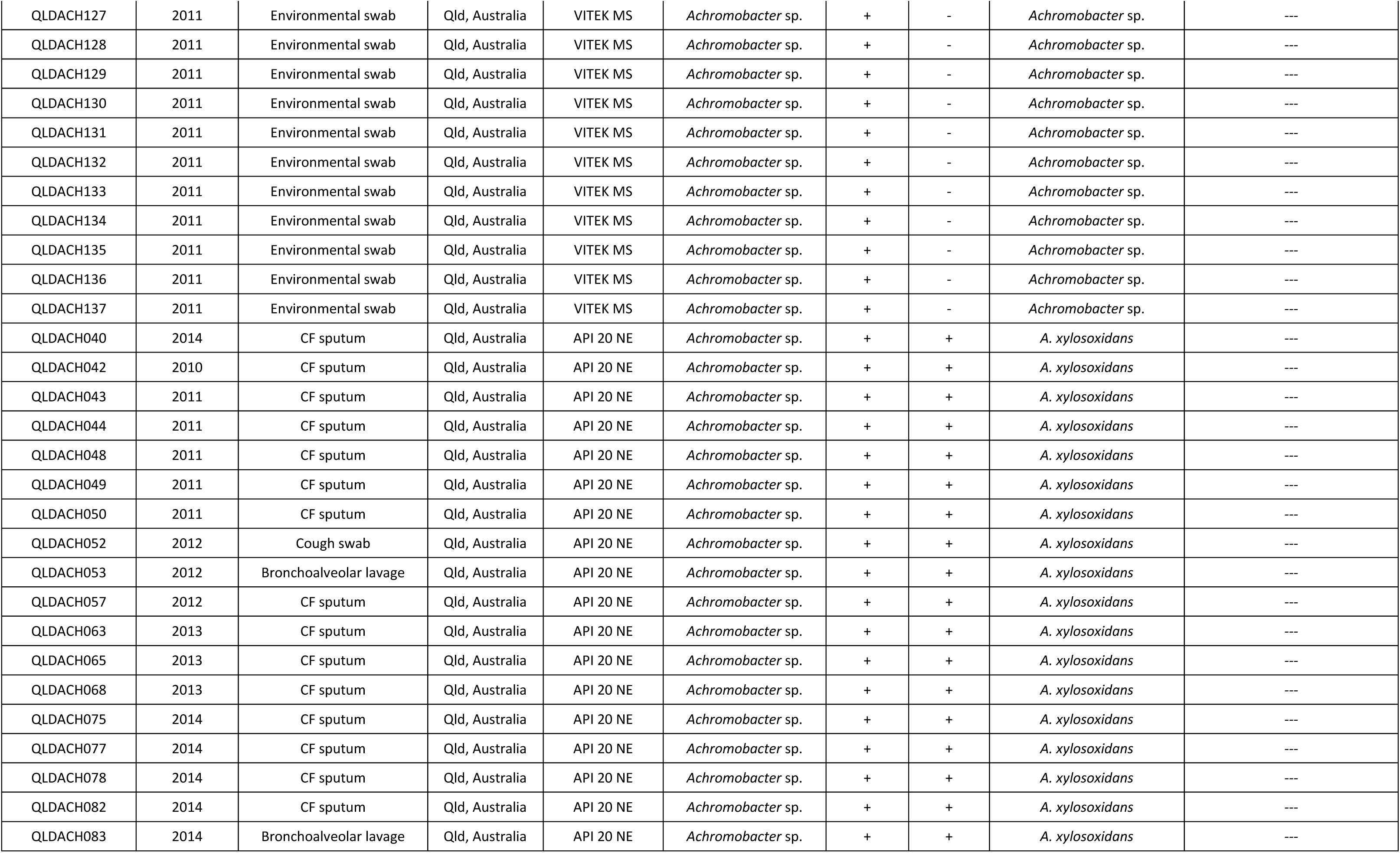
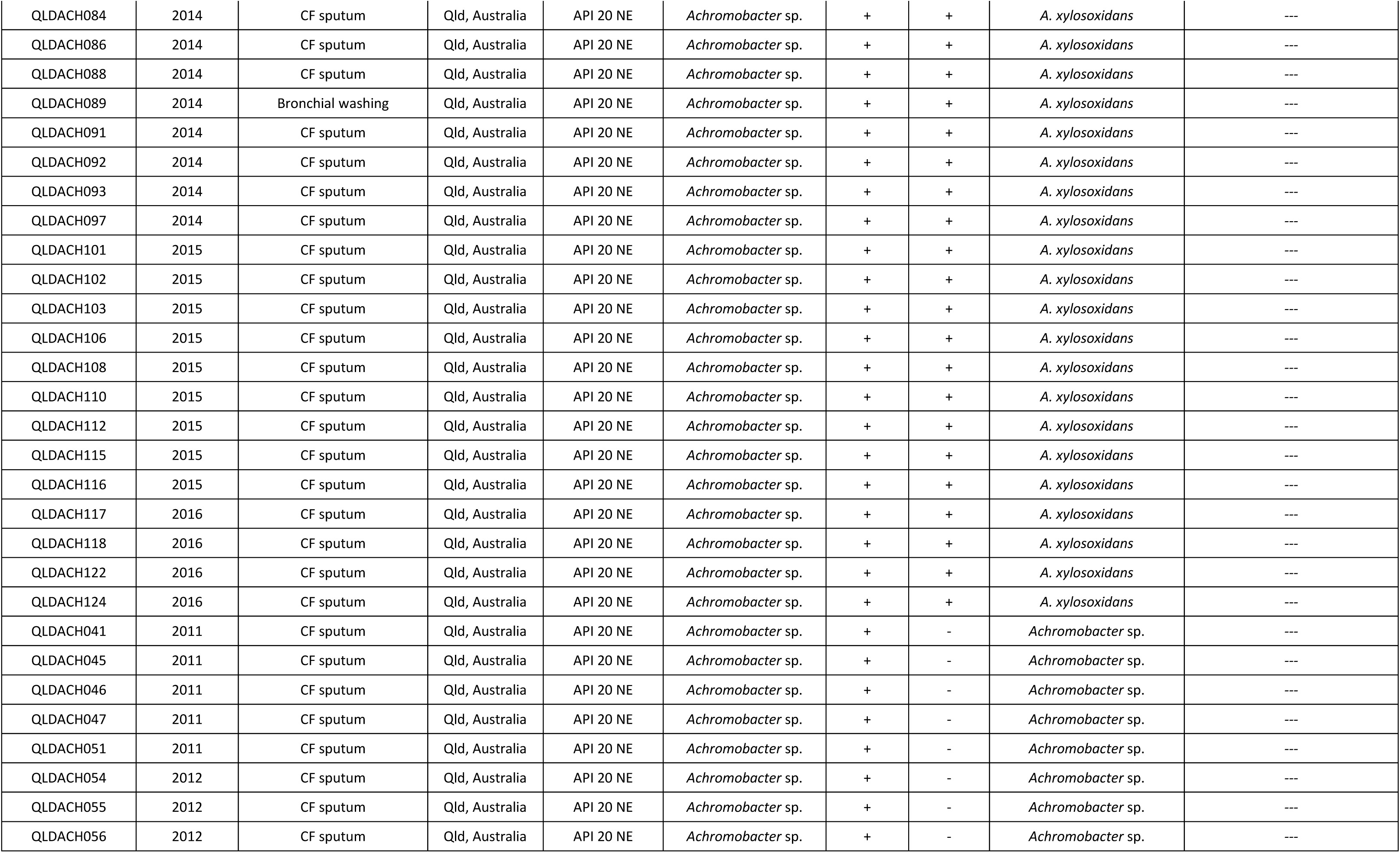
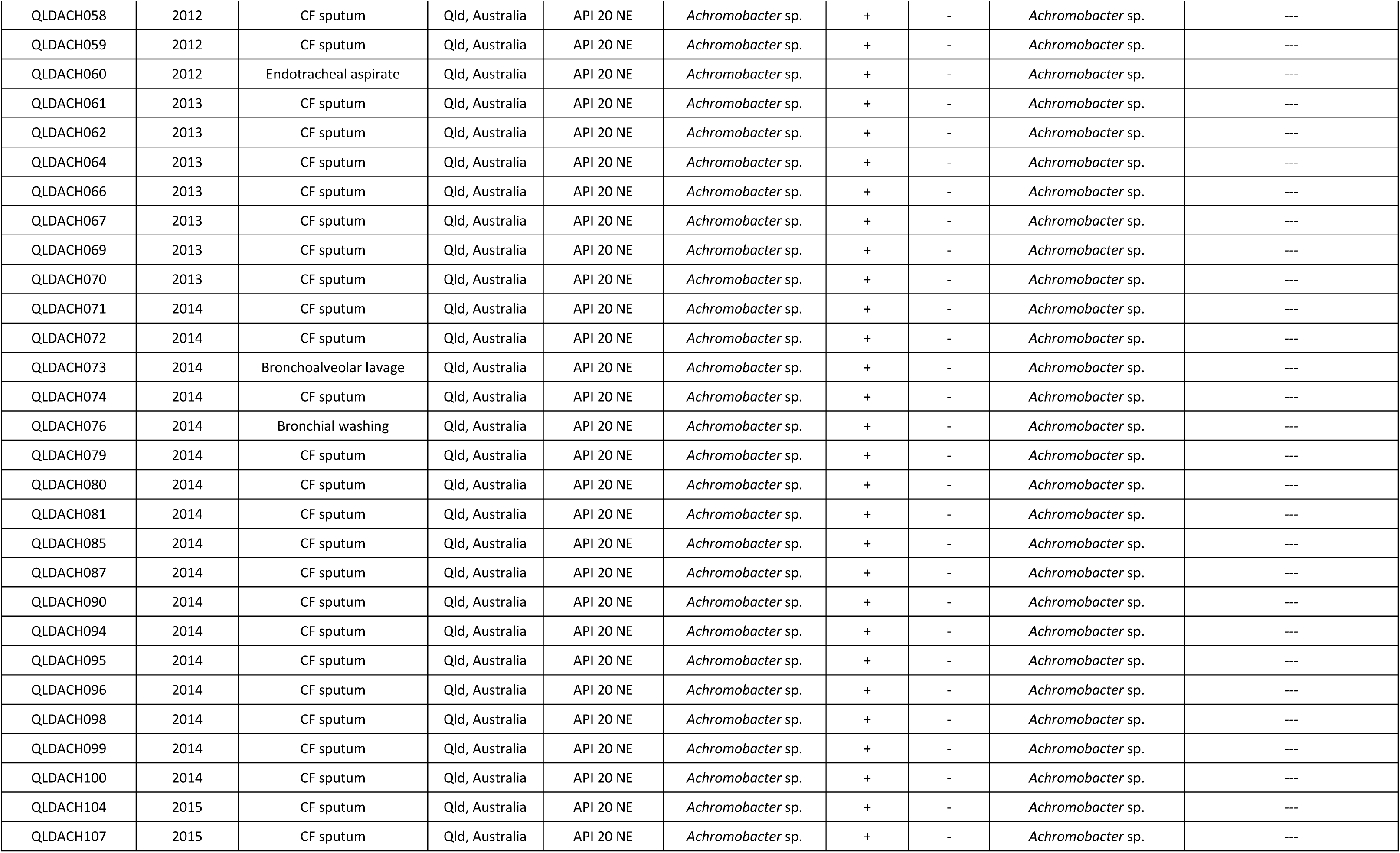
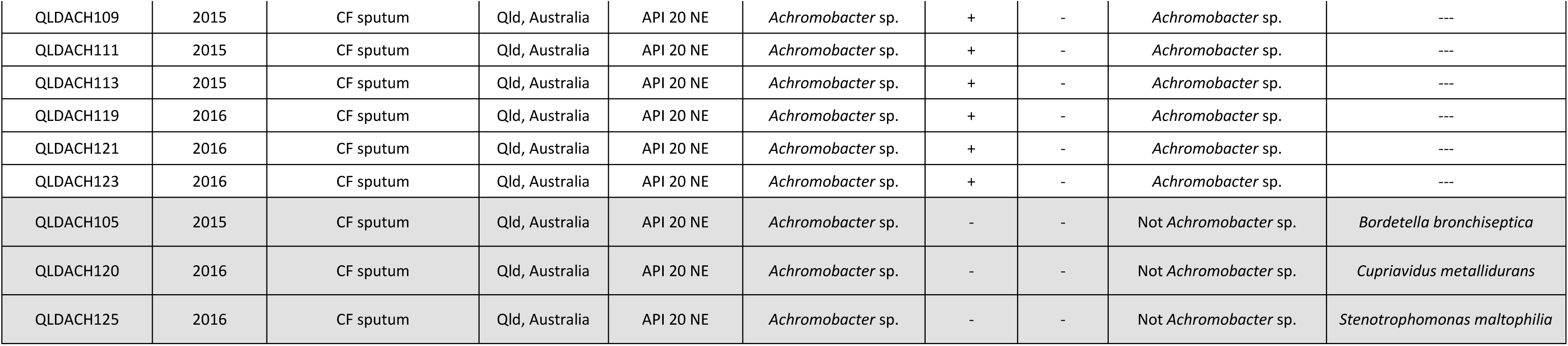
List of the 119 *Achromobacter* isolates examined in this study and associated genotyping/phenotyping results.

